# Pathway and network-based strategies to translate genetic discoveries into effective therapies

**DOI:** 10.1101/051524

**Authors:** Casey S. Greene, Benjamin F. Voight

## Abstract

One way to design a drug is to attempt to phenocopy a genetic variant that is known to have the desired effect. In general, drugs that are supported by genetic associations progress further in the development pipeline. However, the number of associations that are candidates for development into drugs is limited because many associations are in noncoding regions or difficult to target genes. Approaches that overlay information from pathway databases or biological networks can expand the potential target list. In cases where the initial variant is not targetable or there is no variant with the desired effect, this may reveal new means to target a disease. In this review we discuss recent examples in the domain of pathway and network-based drug repositioning from genetic associations. We highlight important caveats and challenges for the field, and we discuss opportunities for further development.

## Introduction

Human genetic association data has revealed how genes influence the susceptibility to rare and complex diseases. A central objective of drug development is to modify disease risk in predictable and beneficial ways, e.g., therapeutic lowering of blood pressure or cholesterol to prevent myocardial infarction. If the direction of perturbation can be inferred from human variation data–e.g., loss-of function, etc.–these ‘natural’ perturbations could be harnessed to indicate a therapeutic hypothesis for rational drug design (1).

In addition, human genetic perturbations may also suggest targets that are the most likely to succeed through drug development. Nelson et al. (2) analyzed the evidence behind drug candidates that entered clinical trials. They found that the fraction of compounds with supporting genetic associations increases throughout the drug development pipeline, suggesting that drugs supported by associations are more likely to successfully advance through the stages of clinical trials where drugs must show efficacy. This highlights the promise of genetic associations that can be mimicked by drugs. In the idealized case, they can take us from a new mechanism to an effective therapeutic.

### The *PCSK9* success story

An illustrative example for this human-genetics informed approach to drug design involves characterization of the Proprotein convertase subtilisin/kexin type 9 (PCSK9) connection to heart disease. In 2003, genetic variation at PCSK9 was determined to be a factor in autosomal dominant hypercholesterolemia (3). Specific variants within the gene - all missense mutations - segregated with the phenotype. However, no nonsense or frameshift mutations were identified, leading to the inference that these variants may be gain of function and that this gain results in higher levels of low-density lipoprotein cholesterol (LDL-C). Subsequently, *PCSK9* was re-sequenced in the extremes of LDL-C, which uncovered nonsense mutations in the gene (4). The authors then designed an assay for two variants and observed that individuals with these alleles exhibited substantially lower LDL levels. In a follow-up study, nonsense mutations in the gene were revealed to be associated with both lower LDL levels as well as a substantial reduction in the lifetime risk of coronary heart disease (5).

Taken together, this genetic data strongly suggested that a therapeutic that mimics the effect of *PCSK9* loss of function should not only lower LDL-C levels but also reduce risk of heart disease. Monoclonal antibodies against *PCSK9* indeed demonstrate the LDL lowering effect in clinical trials (6), and biologics for *PCSK9* are now available and are being tested in phase III clinical trials. Importantly, clinical trials for these agents have demonstrated their ability to lower LDL-C levels even without lifelong exposure to reduced *PCSK9* activity. A recent meta-analysis of existing trials data suggests such treatments will be beneficial (7), and ongoing trials will assess whether or not this translates into a reduction in cardiovascular events after a heart attack (8).

### *PCSK9* is not the only success

The translation of the *PCSK9* genetic discovery to therapeutic is one example among an increasing number of such reports. Missense mutations in the islet-specific zinc transport (*SLC30A8*) are associated with increased risk to type 2 diabetes (9) and loss-of-function mutations are associated with 2-fold protection against diabetes (10). Loss-of-function mutations in the apolipoprotein C3 component (*APOC3*) have been shown to dramatically lower triglycerides, cholesterol levels, and risk of coronary heart disease (11). Loss of function mutations in Niemann–Pick C1-like 1 (*NPC1L1*) are associated with decreased LDL cholesterol and reduced risk of coronary heart disease (12). While mutations in the amyloid precursor protein (*APP*) are known to associated with early-onset Alzheimer’s disease, a recent report identified key mutations which protect against AD and age-related cognitive decline (13). These recently identified examples, analogous to *PCSK9*, offer promising therapeutic hypotheses worthy of further examination.

## From genes to pathways and networks

The *PCSK9* and above examples represent best-case scenarios: a loss of function allele induced a desirable phenotype that could be mimicked by a treatment. This demonstrates the promise of association-based drug development; however, the number of such successes has been limited. The NHGRI GWAS catalog (14) was developed to curate genetic associations and now holds more than 15,000 SNP-trait associations, but we have not yet developed effective drugs based on all of these associations. There have been a limited number of cases where each required elements falls cleanly into place. The success stories have occurred for variants in coding regions, but most associated variants in the NHGRI catalog map to the non-coding genome. For these associations identifying the causal variant and connecting it to the causal gene remains challenging.

Importantly, each variant-phenotype association may not directly suggest druggable target. Pathway and network-based approaches aim to expand the number of associations that lead to effective treatments by expanding the set of potential targets. For example, a genetic association may suggest that altering the effect of a gene would be beneficial, but perhaps that gene’s product can’t be perturbed using biologics or small molecules. A pathway-based strategy could seek to push towards the desired state by targeting genes upstream or downstream in a signaling pathway, while a network-based strategy may seek to target interacting partners to achieve the desired outcome (Figure 1).

**Figure 1:**
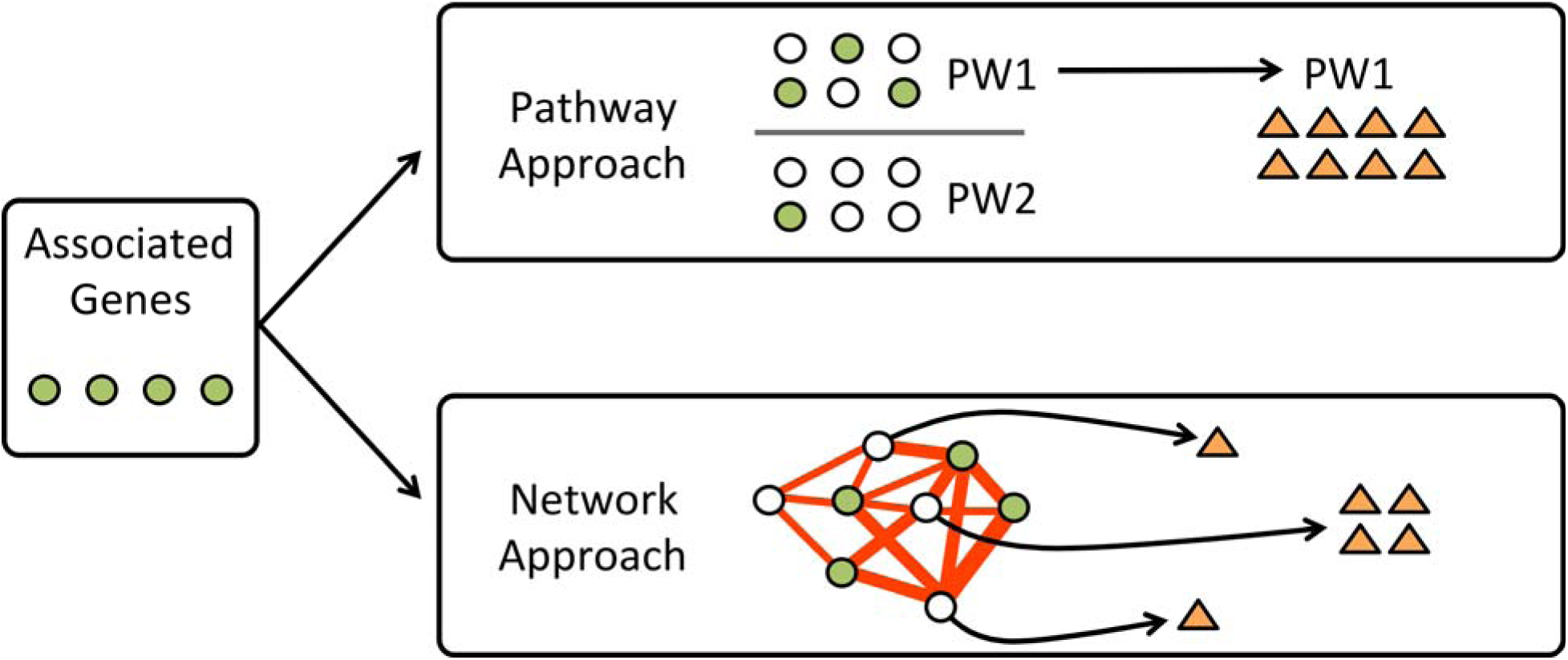
A genetic association study identifies genes associated with a phenotype. If those genes do not directly suggest effective strategies for therapeutics, researchers can use pathway or network-based approaches to expand the potential compound set. In the pathway-based approach, one or more associated pathways are identified and drugs that that target those pathways are prioritized. In the network approach, genes are mapped onto an interaction network and genes that are highly connected to associated genes are identified. Drugs that target those genes represent candidates for further study.

## Pathway and network-based drug identification from genetic associations

Researchers are now developing approaches that analyze genetic associations in the context of networks and pathways to prioritize potential drugs (15–20). There is not yet a standard approach or set of techniques that are widely applied, so we highlight examples of the types of techniques being employed.

Cordell et al. (17) performed a meta-analysis of genetic association data for primary biliary cirrhosis and identified both known and new variants associated with the disease. Pathway-based analysis revealed several associated with the disease, and intersecting these pathways with annotations from the Drug Gene Interaction database (21) highlighted candidates that could modulate pathways associated with the disease. Such analyses lay the groundwork for studies that aim to evaluate the repurposed treatments in models of the disease.

Okada et al. (18) integrated many types of data, in addition to pathway annotations, to identify potential treatments for rheumatoid arthritis (RA). The authors started with a large meta-analysis of genome-wide association data, which provided a set of associated variants. The authors then identified genes that were in associated pathways, showed relevant mouse phenotypes, had protein-protein interaction network support, and many other factors of potential relevance. This allowed the authors to define a score for each gene based on the number of resources supporting its association. They then identified drugs that targeted the top genes. Overlaying drug-target information from DrugBank (22) and the Therapeutic Targets Database (23) revealed that a number of top candidates from the analysis were already targets of existing drugs for RA, suggesting that drugs targeting other top candidates might also be repurposed to treat RA. A similar approach has also been used to identify anti-cancer drugs (20).

Both of the above techniques use curated knowledge to link associations to potential drugs. In contrast with approaches that overlay curated knowledge to calculate a score based on summed sources of evidence, Greene et al. used machine learning methods to identify regions of networks that were informative of genetic associations through a technique termed NetWAS (16). The NetWAS method uses the results of a genome-wide association study to find these regions of the networks. The analysis described in the paper used genes that had a nominally significant (uncorrected p < 0.01) association with hypertension endpoints. Genes identified by the network analysis were more likely to be associated with hypertension-related gene ontology processes (24) and more likely to have been annotated as hypertension genes in the OMIM database (25). These genes were also more likely to be the target of antihypertensive therapies in the DrugBank database (22), suggesting that compounds that target other highly-ranked genes would be candidate antihypertensive treatments.

The highlighted techniques focus on drug repurposing for a single disease starting from a single genome-wide study or meta-analysis. In contrast with these, Himmelstein et al. (19) are carrying out an effort to perform large-scale drug repurposing from a catalog of genetic associations. They use heterogeneous networks that connect genes, diseases, tissues, and other factors (26). They then apply machine learning methods to predict effective drug-disease pairs. The entire process has been visible on an open science platform called Thinklab. The open platform reveals the extensive sets of analyses that lay the groundwork for machine learning applications. Though work is ongoing the initial results are highly promising.

## Challenges for network and pathway-based approaches

While promising results of network and pathway-based drug discovery approaches are emerging, methods must still overcome some challenges. A key step for all network and pathway-based approaches is to connect associated variants to the relevant gene or genes that are causal for the phenotype of interest. Methods developers need to take into account potential sources of bias in how the underlying SNPs are mapped to a candidate gene. For example, genes with more SNPs are often more central in many networks and more annotated in pathway resources (27, 28). Network and pathway-based approaches need to use analytical strategies that map SNP-based p-values to gene-based p-values without suffering inflation from large genes (29, 30). Approaches that fail to account for this structure may simply identify generic, as opposed to phenotype-specific, associations.

In addition to such biases, there are important biological challenges to mapping variants to genes. A variant may affect a distal, as opposed to the closest, gene. An recent example which highlights this issue is intronic variation near an obesity-associated locus (31–33). It is now clear that this is due, at least in part, to long range interactions between these variants and the genes *IRX3* and *IRX5* (34, 35). The variant to gene mapping problem remains a challenging area of work. Techniques to predict enhancer-gene interactions (36) or those that use expression quantitative trait loci (37) may help to improve accurate variant to gene mappings. This is an active area of research, and we expect improvements in this area to immediately improve the sensitivity and specificity of pathway-based methods.

An often-overlooked aspect of studies that incorporate multiple data types and sources for drug repurposing is licensing and resource accessibility. The manner in which restrictive database licenses have hampered progress by the Himmelstein et al. team (19) has been instructive (38). For example, the authors have been unable to use the MSigDB database of chemical and genetic perturbations (39). While many resources do allow reuse by academic users, unclear and restrictive licenses have the potential to sap time from researchers engaged in network and pathway-based drug repurposing studies. This demonstrates that practitioners need to consider not only whether or not resources may provide useful data but also whether or not using such resources in a robust and reproducible way is compatible with their licenses.

## Opportunities on the horizon

Network and pathway approaches benefit from the ability to use multiple distinct associations to identify a relevant pathway or region of a biological network. While individual phenotypes have provided a fruitful starting point, collections of phenotypes may be an even more promising avenue for such approaches. We might expect that related phenotypes may not be influenced by the same variant and genetic mechanism, but the same pathway could drive them. An early report suggests that the use of such multi-phenotype or phenome-wide data for drug repositioning may be fruitful (40).

The Alzheimer’s Disease Neuroimaging Initiative has collected and shared genotype, fluid biomarker, and imaging data on a set of individuals (41). These data can be used for network and pathway-based association analysis of individual phenotypes (42). Multi-phenotype analysis may identify consistent pathways or network regions associated with certain imaging phenotypes, which could suggest potential interventions.

Another major source of phenotyping data is the electronic health record (43, 44). The breadth of these data presents both challenges and opportunities (45). Because these records contain numerous phenotypes with treatments available, existing drug-disease pairs can be used to evaluate methods.

In a few cases, a genetic association may directly suggest a drug-development strategy. For most cases, translating genetic associations into treatments is likely to be complex. In these cases, network and pathway-based approaches can integrate information from multiple types of variants with distinct effects to reveal pathways that can be modulated to affect a phenotype. The expanding availability of sequence measurements with extensive phenotyping makes this a promising area for the development of new methods. Systems capable of converting such data into drug candidates may help researchers repurpose existing therapies for other uses and target drugs for development more efficiently.

